# Decoding synonymous codon selection with a Transformer model

**DOI:** 10.64898/2026.03.28.714798

**Authors:** Hélène Bret, Ingemar André

**Affiliations:** Lund University, Lund, Sweden

**Keywords:** codon choice, rare codons, transformers, language models

## Abstract

The genetic code is highly redundant, with many synonymous codons encoding the same amino acid. Codon usage influences RNA structure, signaling, and translation rates. Differences in tRNA availability modulate elongation, with rare codons slowing translation and affecting co-translational folding and gene expression. Despite their functional importance and non-random distribution, rare codons are underrepresented in natural datasets, restricting the development of predictive models. We developed a transformer-based model that predicts codon sequences from amino acids, substantially improving rare codon prediction. The model learns codon signatures encoding species identity, RNA thermodynamic properties, and elongation constraints without explicit labels. Attention analysis shows that codon choice depends on both short and long-range sequence contexts, recovering dicodon effects and highlighting additional motifs. Finally, predictions correlate with experimental measurements of the impact of synonymous mutations on protein fitness, linking gene sequence to fitness and functional consequences, providing a framework to connect sequence variation, translation, and protein function.

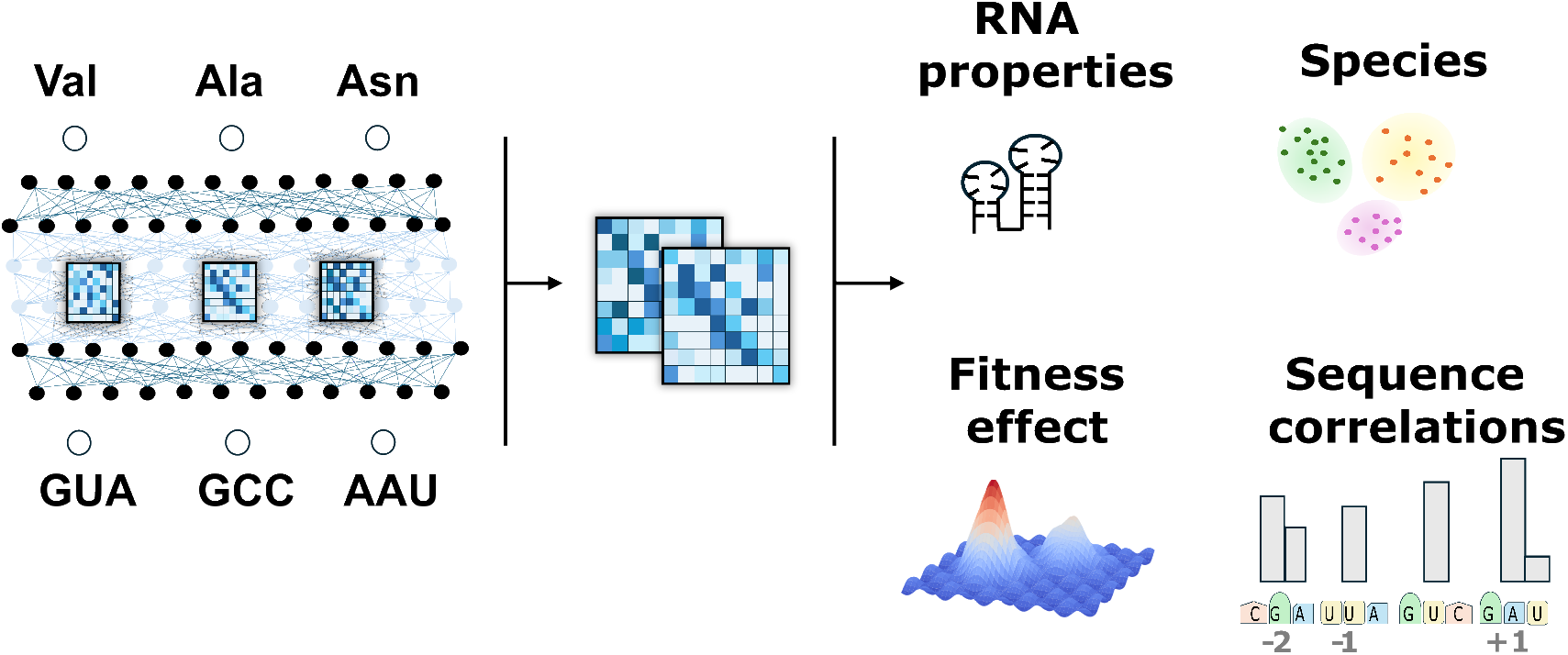

## 1 Introduction

The genetic code is redundant: most amino acids are encoded by multiple synonymous codons, ranging from one to six. These synonymous codons differ in their nucleotide composition, most often at the third position of the triplet. Such variations are increasingly recognized as important factors influencing cellular processes. They can affect mRNA secondary structure and stability^1–3^, and even modulate the expression of nearby genes^4^, thereby impacting translational efficiency and RNA turnover. Recent studies also suggest that the usage of synonymous codons in humans may contribute to protecting against undesirable RNA species^5^, such as aberrant splicing products or viral transcripts. Beyond RNA-level effects, synonymous substitutions can modify cis-regulatory elements within coding regions^6^, including exonic splicing enhancers^6,7^, microRNA binding sites^8–10^, and translational pausing motifs^11,12^.

Importantly, synonymous codons are not used uniformly: their decoding depends on the abundance and availability of cognate tRNAs, which vary across species and tissues^13,14^. This differential tRNA supply can modulate translation kinetics, especially during co-translational protein folding, by creating local pauses that coordinate domain folding or protein–protein interactions^15–17^. Codons associated with low tRNA abundance, often termed rare or slow codons, can therefore act as regulatory elements that shape the temporal and structural dynamics of translation. Controlled slowing of translation has been shown to be crucial for efficient folding, subunit assembly, and post-translational targeting of many proteins^11,18,19^.

To quantify and model these patterns, several statistical indices have been developed, such as the Codon Adaptation Index (CAI; ^20^), the tRNA Adaptation Index^21^, and related measures like MMinMax^22^ or Relative Synonymous Codon Usage (RSCU) ^23^. These approaches summarize codon usage bias or translational efficiency at the gene level and remain valuable descriptors of global codon preferences. However, they are limited in capturing context-dependent or sequence-level determinants of codon choice, such as local structural or compositional features that influence rare codon placement.

More recently, machine learning and deep learning approaches have been developed to learn codon choice directly from sequence data^24,25^. Among these, Transformer-based architectures have shown particular promise due to their ability to model long-range dependencies in biological sequences^26,27^. Still, most existing models are designed for codon optimization, aiming to maximize protein expression in heterologous systems^28–30^. Such models typically operate at the nucleotide level, taking a reference coding sequence as input and modifying it toward host-specific codon preferences^24,25^. Moreover, they are often trained on limited taxonomic scopes, restricting their capacity to capture the broader evolutionary landscape of synonymous codon usage.

Recent Transformer-based models such as CodonTransformer^27^ represent a major step toward sequence-level prediction of synonymous codon usage. However, these architectures mainly focus on overall accuracy and tend to reproduce the codon biases present in their training data, rather than explicitly capturing the factors that determine rare codon placement. As a result, they often miss the subtle context-dependent patterns of rare codon usage, which are increasingly recognized as important for co-translational folding, translation dynamics, and protein expression regulation..

Here, we introduce CaNAT (Codon from Amino Acid with a Non-Autoregressive Transformer), a deep learning model designed to predict synonymous codon usage directly from protein sequences (Fig. 1 A). In contrast to previous approaches, CaNAT was trained with a balanced codon weighting strategy, ensuring that rare synonymous codons contribute equally during optimization. This design allows the model to better capture the contextual determinants of rare codon selection, beyond global frequency bias. Trained on a large and taxonomically diverse dataset comprising more than three million coding sequences from more than 600 species, CaNAT generalizes across evolutionary and compositional contexts. The model predicts the most probable native codon sequence for a given amino acid input, providing both accurate reconstruction of natural coding sequences and interpretability of the factors influencing codon choice. This framework bridges sequence modeling and codon-bias analysis, offering new opportunities to investigate how local sequence context and evolutionary constraints shape synonymous codon usage.

**Figure 1:**
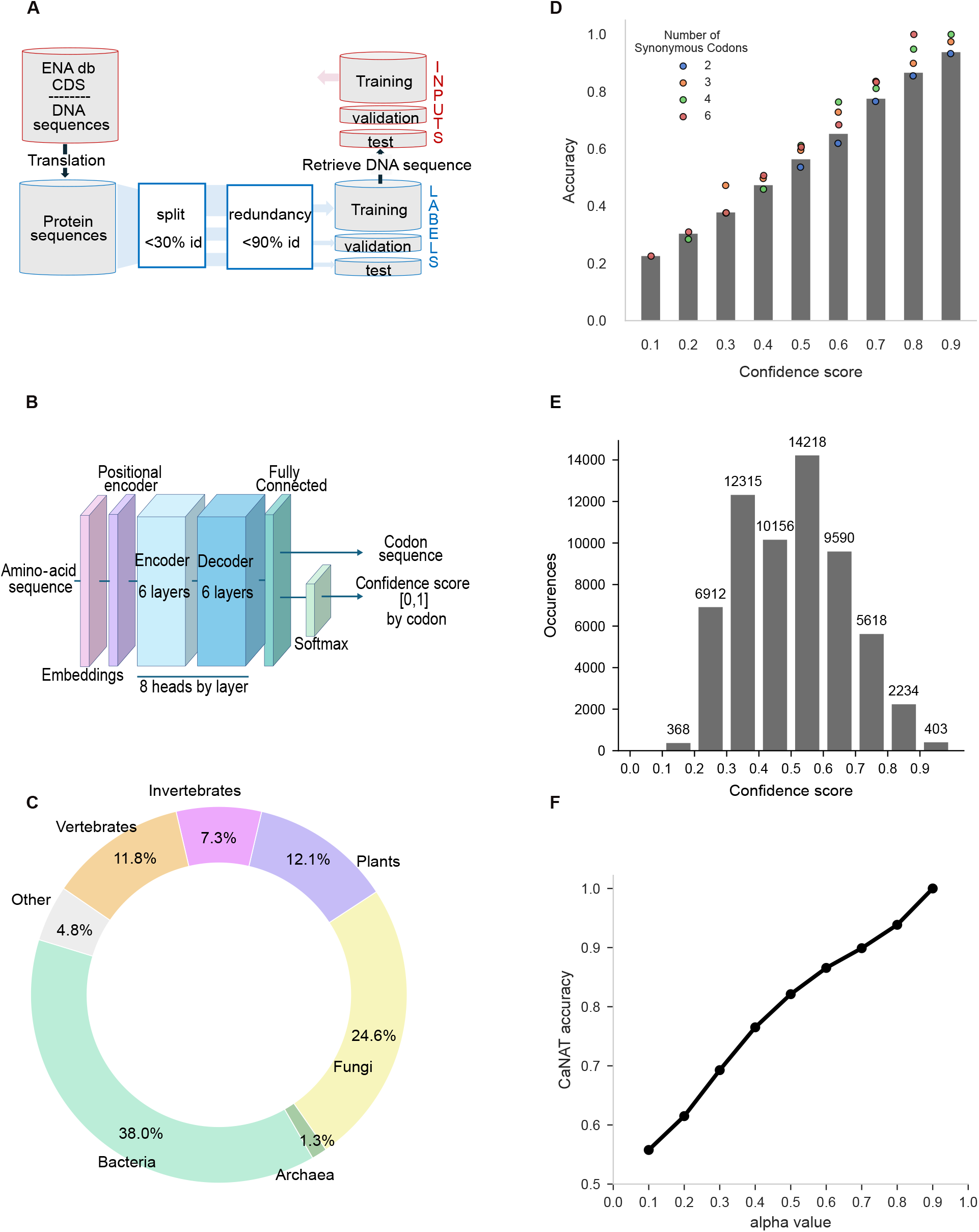
Model architecture, training data, and confidence–accuracy relationship. (A) Pipeline of independant construction of training, validation, and test datasets (B) CaNAT architecture predicting codon sequences from amino acid sequences and outputting a confidence score. (C) Taxonomic composition of the training set across major taxonomic groups. (D) Prediction accuracy on the test set as a function of confidence (bins of 0.1). Colored points indicate accuracies for each number of synonymous codons, which are not directly comparable. (E) Distribution of confidence values for all codon predictions in the test set (Detail per number of synonymous codon in Fig. S1). (F) Relationship between normalized confidence level 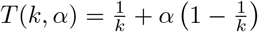, where k is the number of synonymous codons, and prediction accuracy, enabling comparison across codon degeneracy levels.

## 2. Results

### 2.1 CaNAT provides reliable confidence scores for codon prediction

We developed a non-autoregressive Transformer model, CaNAT, designed to predict full codon sequences directly from amino acid inputs without masking(Fig. 1 A,B). Preliminary tests showed that masking strategies did not provide measurable benefits, so training was performed without them. CaNAT was first trained on synthetic sequences to capture the genetic code, followed by large-scale training on natural coding sequences.

A major challenge in this task arises from the strong bias in synonymous codon usage. Because sequences must remain biologically valid, the training set could not be fully balanced. To compensate, we implemented a batch-wise weighted cross-entropy, ensuring that gradients were appropriately scaled during optimization and preventing the underrepresentation of rare codons.

The training set was curated to avoid overrepresentation of any gene family and maximize taxonomic diversity, ultimately comprising 620 species and over 3 million filtered coding sequences from European Nucleotide Archive (ENA) cRNA (Fig. 1 A,C) (See sectionMethods). Separate validation and test sets were constructed for an unbiased evaluation. CaNAT includes 6 encoder and 6 decoder layers, each with 8 attention heads and 512-dimensional embeddings, allowing all codons to be predicted simultaneously rather than sequentially, substantially accelerating both training and inference (Fig.1 B).

CaNAT generates two outputs: a predicted codon sequence (excluding the stop codon, which was removed due to limited training examples) and a codon-level confidence score ranging from 0 (low) to 1 (high) (Fig.1 E, detailed by number of synonymous codons Fig. S1). The codon-level confidence score is defined as the predicted probability of each codon after applying a softmax over all codons and correlates strongly with prediction accuracy (Fig. 1 D), indicating that the model effectively measures its own uncertainty (“knows when it knows”). Confidence levels depend on the number of synonymous codons per amino acid (Fig.1 D); for example, a medium confidence score among six codon choices already reflects high certainty compared to a binary alternative. To enable codon-wise comparisons, thresholds were adjusted according to the degeneracy k of each position. Thresholds were calculated using

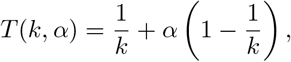

where *α* is a tunable parameter representing the confidence level: *α* = 0 corresponds to a threshold equal to the uniform expectation 1*/k* (i.e., no confidence beyond random choice), and *α* = 1 corresponds to a perfect prediction. Applying this degeneracy-adjusted threshold, increasing the confidence level *α* improves accuracy by selecting positions that are better predicted. (Fig. 1 F).

### 2.2 CaNAT outperforms existing codon prediction models, particularly for rare codons

CaNAT performance was evaluated on an independent test set and compared with several statistical baselines. The first baseline corresponded to an optimal codon choice, defined as selecting for each amino acid the most frequent codon observed in the training dataset, yielding an accuracy of approximately 48%. A second baseline reproduced the natural codon distribution by sampling synonymous codons according to their observed frequencies in the dataset, achieving an accuracy of about 39%. As a lower bound, a random synonymous codon assignment at each position was considered, resulting in an accuracy of 33%. CaNAT reached an accuracy of 53%, exceeding the performance of all statistical baselines (Fig 2 A).

**Figure 2:**
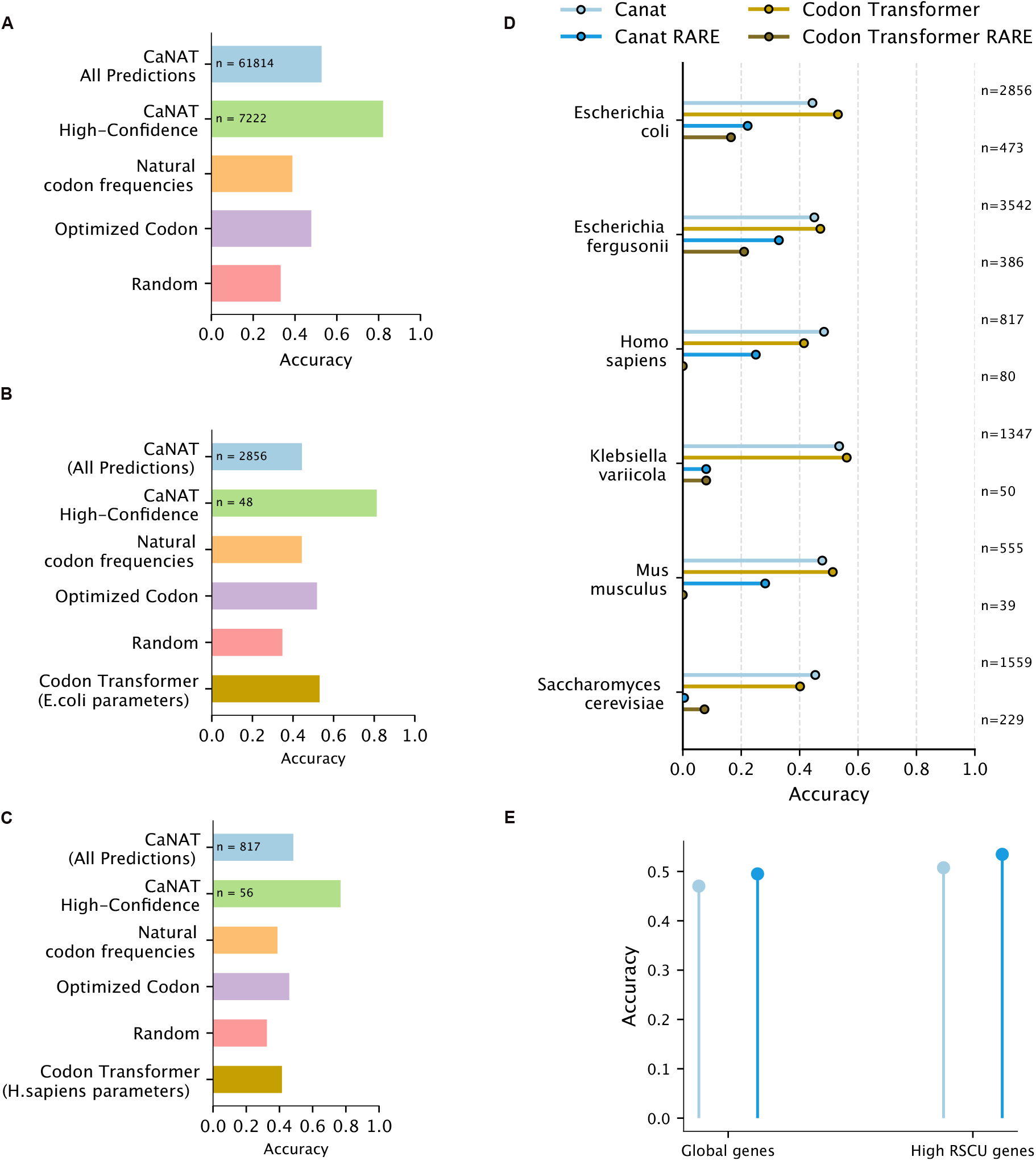
CaNAT performance compared with statistical baselines and species-specific CodonTransformer. (A) Prediction accuracy on the full test set across all species. Bars show the accuracy of the model on all codons, on codons predicted, and for three statistical baselines: optimal codon (most frequent synonymous codon in the test set), frequency-based sampling, and uniform sampling. (B) Same analysis as in (A) restricted to Escherichia coli sequences, including comparison with the species-specific CodonTransformer model using Escherichia coli parameters. (C) Same analysis as in (B) for Homo sapiens, using CodonTransformer parameters optimized for Homo sapiens. (D) Accuracy on codons from species shared between the test set and CodonTransformer training sets, shown for all codons and rare codons (RSCU < 0.7), for both models. (E) Focus on highly expressed E. coli genes (genes with mean RSCU values exceeding the 90th percentile of all genes): accuracy for all codons and rare codons across all genes and within the optimized subset.

To benchmark CaNAT against existing approaches, we compared its performance with CodonTransformer^27^, a recent amino acid–to–codon prediction model fine-tuned separately for each organism. We evaluated both methods on test-set sequences from two model species, *Escherichia coli* and *Homo sapiens*, while retaining the same statistical baselines described above (Fig 2 B–C). Remarkably, on *H. sapiens* sequences, CaNAT outperformed all other methods, including CodonTransformer and the organism-specific optimal codon baseline (the most frequent codon for each amino acid in *H. sapiens*). Conversely, on *E. coli* sequences, CodonTransformer and the optimal codon baseline performed slightly better than CaNAT. However, when restricting the analysis to positions with confidence scores exceeding the adaptive threshold defined at a confidence level of *α* = 0.5 (i. e. k=2: T = 0.75, k=3, T=0.667, k=4: T=0.625, k=6: T=0.583), CaNAT consistently achieved higher accuracy than any other methods (2 A-C). These results indicate that while transformer-based models may be sensitive to species-specific codon biases, CaNAT’s confidence-aware predictions enable targeted performance improvements across organisms. Interestingly, accuracy measured on the training set (2 E) of *E. coli* show a similar accuracy level, meaning that the model is not overfitted.

Considering the subset of species common to both the CodonTransformer fine-tuning datasets and the CaNAT test set (*Escherichia Coli, Escherichia fergusonii, Homo Sapiens, Klebsiella variicola, Mus musculus, Saccharomyces cerevisiae*), CodonTransformer achieves slightly higher accuracy when all codon positions are considered in four of the six species evaluated. However, CANAT clearly outperforms CodonTransformer when focusing on rare codons, defined as those with an RSCU value below 0.7 according to the Kazusa codon usage tables^31^ (See sectionMethods) (Fig. 2D), for all species except *Saccharomyces cerevisiae*, where rare codon prediction accuracy is low for both models. This difference is particularly pronounced in *Homo sapiens* and *Mus musculus*. A likely explanation is that bacteria represent 142 of the 164 organisms in CodonTransformer’s training dataset prior to species-specific fine-tuning; despite fine-tuning, residual biases from bacterial codon usage may persist and negatively impact detection of rare codon context in eukaryotic genomes.

This observation suggests that although CaNAT may be less tuned for globally frequent codons, it captures with greater precision the subtle sequence determinants underlying the placement of rare codons. Since these rare codons often occur at positions that are functionally or structurally important, such as regions influencing translation kinetics or co-translational folding, the ability to predict them accurately may reflect a deeper understanding of biologically meaningful patterns rather than a simple statistical fit to codon frequencies.

Rare codons are unevenly distributed across genes. Highly expressed or housekeeping genes, which are optimized for rapid translation, contain fewer rare codons^32^. Consequently, a rare codon in these genes is more unexpected, highlighting its potential functional importance. Notably, prediction accuracy for these rare codons remains comparable to that across all genes, indicating that CaNAT does not rely solely on global gene expression cues but also captures local sequence patterns to inform codon choice (Fig. 2 E).

### 2.3 Organism-specific codon usage biases emerge naturally from amino-acid sequences

Codon usage preferences are known to be highly organism-specific and represent an essential determinant for accurate sequence prediction. CaNAT was trained without any information regarding the source organism and was not fine-tuned on a per-species basis. To evaluate whether the model nevertheless captures organism-level codon usage biases intrinsically, we compared predicted and empirical synonymous codon distributions for each amino acid across multiple representative species Fig. S2) even for amino acids encoded by up to six codons (Fig. 3 A). For both *E. coli* and *H. sapiens*, the predicted distributions closely recapitulated natural codon usage patterns, with a median Spearman correlation close to 1 across amino acids, indicating that CaNAT implicitly learned species-dependent biases in codon selection. This agreement extended to an extremophile organism, *S. thermophilus*, exhibiting a strong codon usage bias and limited representation in the training dataset, suggesting that the model generalizes such preferences even under data-sparse conditions.(Fig. 3 B,C) Notably, rare codons were preserved in the predicted distributions rather than being replaced by more frequent alternatives, further supporting the model’s capacity to reproduce biologically realistic codon usage.

**Figure 3:**
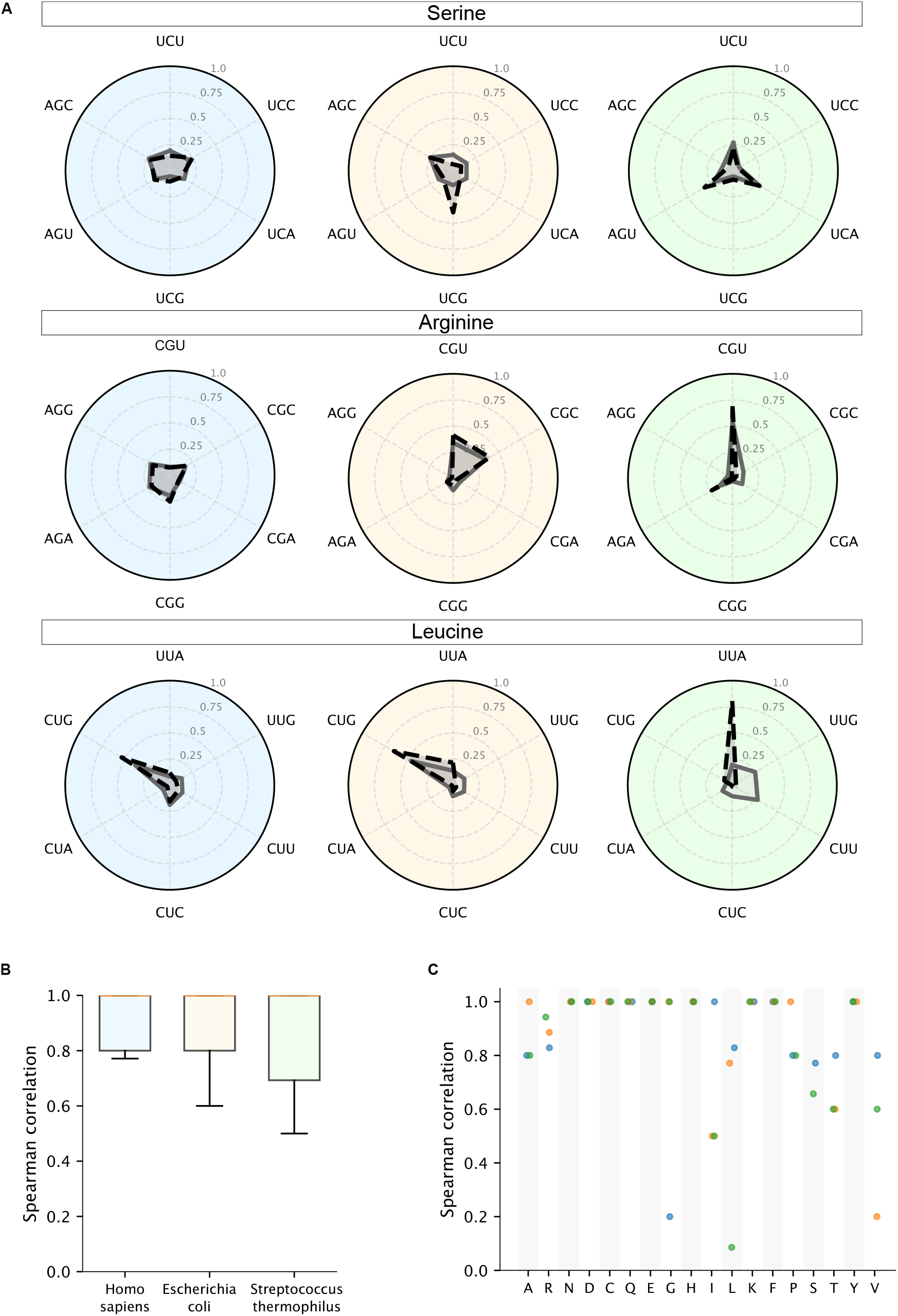
Synonymous codon usage and model predictions across species. (A) Radar plots showing the distribution of synonymous codons for three amino acids (R, L, S) in three species (H. sapiens, E. coli, S. thermophilus). Solid lines indicate the observed codon distributions in the test set, dashed lines indicate model-predicted distributions. (B) Distribution of amino-acid–level Spearman correlations for the three species (C) Spearman correlation between observed and predicted codon distributions for each amino acid, calculated separately for each species (blue: *H*.*sapiens*, orange: *E*.*coli*, green: *S*.*thermophilus*).

CaNAT also maintained the rank order of synonymous codons, accurately reflecting their relative frequencies for the majority of amino-acid, even displaying markedly skewed usage (Fig. S3-S6). This observation suggests that the occurrence of rare codons in CaNAT predictions does not arise from stochastic variability among synonymous codons but from specific contextual signals captured by the model. Although minor deviations were observed, they fall within the expected range of variation for learned distributions, confirming that CaNAT models codon usage trends faithfully rather than relying on memorization of global frequencies.

These results show that CaNAT does not merely reproduce overall codon frequencies but instead captures the intrinsic statistical structure of organism-level codon usage. The model appears to encode global tendencies in synonymous codon selection that are consistent with species-specific translation patterns. The preservation of rare codons in its predictions further suggests that CaNAT has internalized subtle evolutionary or functional constraints, enabling it to reproduce realistic codon usage distributions across diverse organisms.

To further investigate how organism-level information is represented within the model, we analyzed the embeddings produced by CaNAT at each codon position using kernel density estimation and linear discriminant analysis (LDA) (Fig. 4). Remarkably, embeddings clustered by organism even when analyzed at the positional level rather than across entire sequences, indicating that species-specific signatures are encoded consistently throughout the sequence. This positional resolution suggests that CaNAT has internalized organism-dependent codon usage patterns deeply enough for the identity of the source organism to be inferred directly from local embedding features.

**Figure 4:**
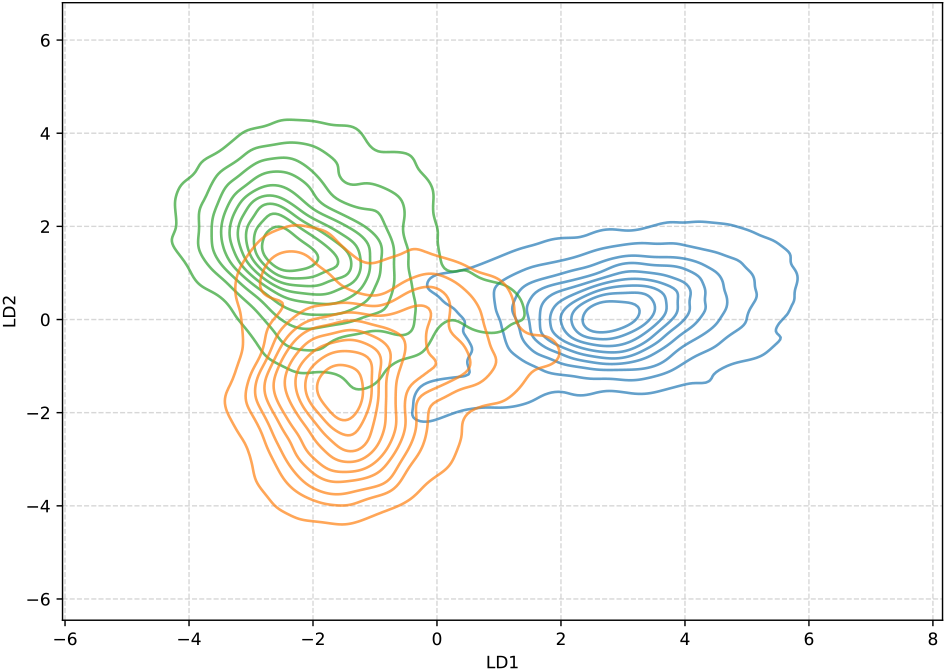
Species separation in CaNAT embedding space. Linear discriminant analysis (LDA) of model embeddings, with contour representation highlighting separation between *H*.*sapiens* (blue) *E. coli* (orange) and *S. thermophilus* (green).

### 2.4 Biological constraints shaping codon choice are intrinsically reflected in CaNAT internal representations

Multiple factors are known to influence codon choice, including nucleotide composition, RNA secondary structure, and local sequence context. By analyzing CaNAT accuracy determinant and attention maps, we investigate how these biological constraints are represented in the internal features used for codon-level prediction and how they relate to synonymous codon choice.

RNA stability has been proposed as an important determinant of codon usage. We therefore asked whether prediction accuracy is modulated by RNA stability.

The relationship between RNA sequence stability and prediction accuracy was evaluated while accounting for GC-content, a species-dependent factor that influences RNA stability through nucleotide composition. Two linear models were fitted: one using GC-content alone to predict accuracy, and a second including both RNA stability and GC-content. Incorporating RNA stability increased the explained variance from 15% to 19% (R^2^; Tab. 1) Fig. S7, indicating that stability provides additional information on prediction accuracy beyond GC-content. These results suggest that CaNAT captures aspects of RNA stability, including effects from secondary and tertiary structure, that contribute independently to codon prediction accuracy.

**Table 1:** GC-content and stability relevance in accuracy explanation. Based on a random sample of 10,000 sequences from the training set.

|  | R<sup>2</sup> | p-value |
| --- | --- | --- |
| GC-content | 0.148 | $p < 10^{-3}$ |
| GC-content + stability | 0.191 | $p < 10^{-3}$ |

In addition to capturing biophysical and organism-level features, several attention heads exhibit distinct diagonal patterns over the sequence (Fig. 5 A). Such diagonal patterns are observed in 16 attention heads spread across 4 Transformer layers. This suggests that 16 of the 96 total attention heads are dedicated to pairs of sequence-position–related motifs, while the remaining heads capture more diffuse information that is not easily recognizable visually (Fig. S8).

**Figure 5:**
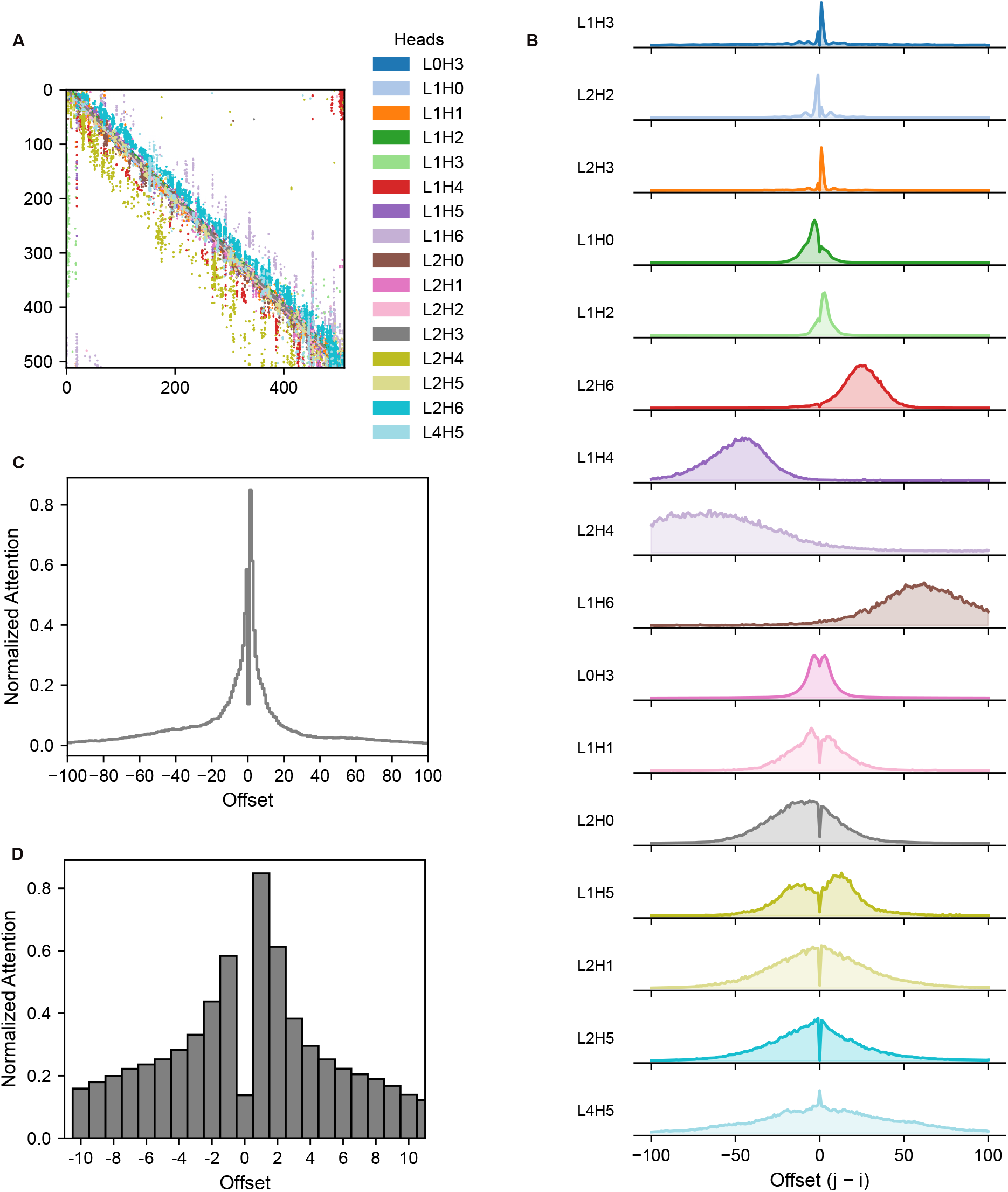
Visualization of model attention patterns. (A) Contours of averaged and normalized attention maps across all test-set sequences. Each color represents a different attention map, highlighting distinct model focus areas. (B) Ridge plots of attention distributions for each attention map as a function of sequence offset (−100 to +100), summed across all test-set sequences. Colors correspond to those in panel A. (C) Histogramm of mean normalized attention values across all attention maps, zoomed to offsets −100 to +100, emphasizing fine-scale attention patterns around central positions. (D) Histogramm of mean normalized attention values across all attention maps, zoomed to offsets −10 to +10, showing a peak of attention on position +1 and a bias toward downstream positions

**Figure 6:**
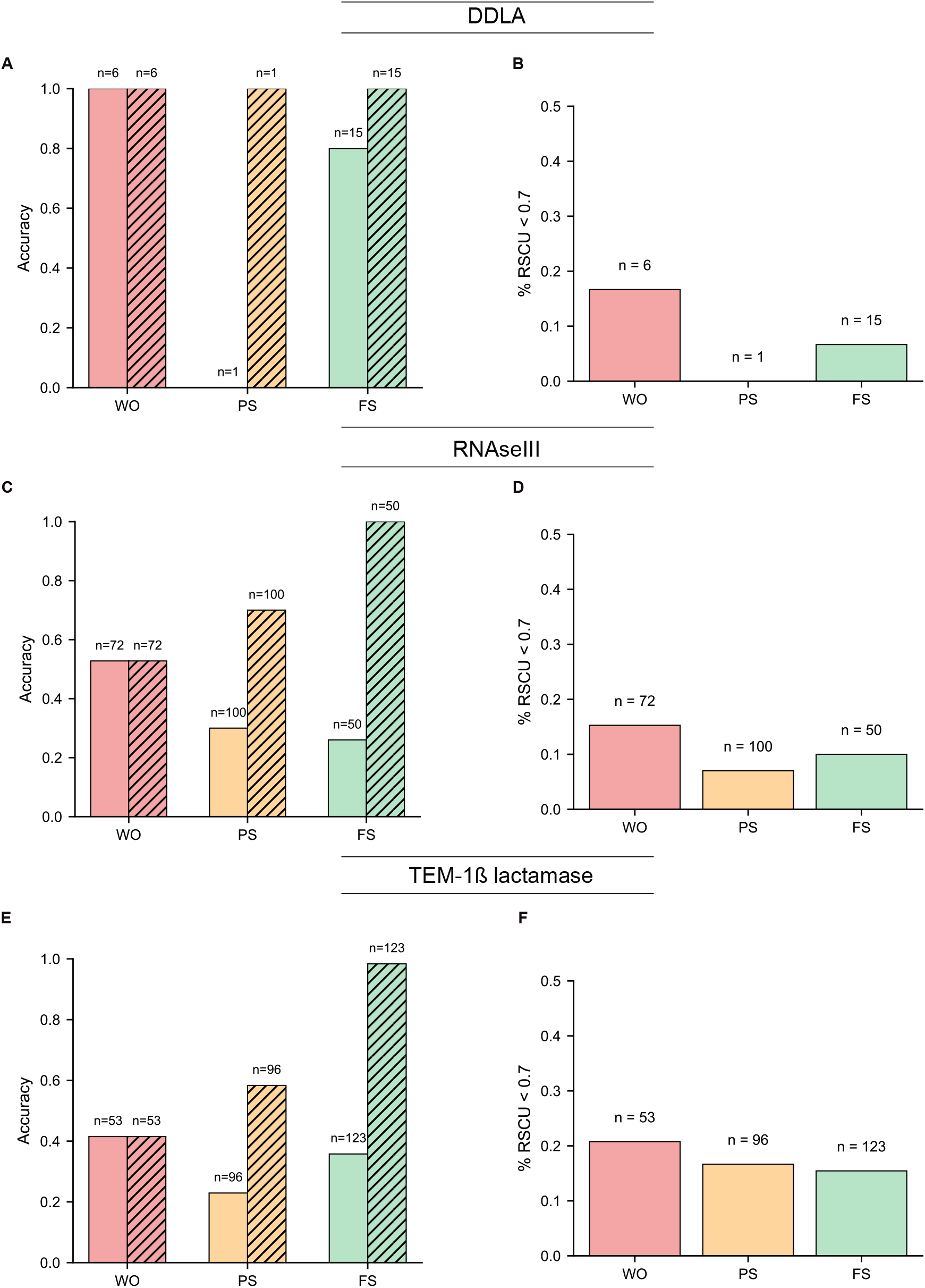
Prediction of fitness effects of codons by CaNAT. (A) Accuracy of predictions on ddlA for positions classified as Wild Type only (WO), Partially Synonymous tolerant (PS), and Fully Synonymous tolerant (FS), shown against the Wild Type codon (solid lines) and all tolerated codons (dashed lines). (B) Percentage of codons with RSCU 0.7 at WO, PS, and FS positions. (C-D) Same as (A-B) for RNase III. (E-F) Same as (A-B) for TEM-1 *β*-lactamase.

These diagonal attention patterns can be grouped into three main categories (Fig.5 B). The first category (L1H3 (Layer 1 Head 3), L1H0, L2H2, L2H3, L1H2) comprises tight diagonals located very close to the main diagonal, corresponding to interactions between nearby sequence positions (small offsets). In these heads, attention is strongly concentrated on a limited number of adjacent positions, forming a well-defined peak. This pattern is consistent with dicodon (codon-pair) effects, whereby adjacent codons are not used independently^33,34^. Instead, specific codon pairs are preferentially favored or disfavored in a directional manner, depending on local sequence context and underlying dinucleotide biases^35^ and influences by itself RNA properties and translation elongation^36–38^.

A second category (L2H6, L1H4, L2H4, L1H6) corresponds to long-range attention patterns, characterized by diagonals located far from the central diagonal. These heads capture interactions between distant sequence positions, with offsets typically ranging from approximately − 70 to +60. Such patterns suggest that the model incorporates long-distance contextual information when predicting synonymous codons. The broader attention window indicates that this does not correspond to a precise pair of positions but rather to a broader motif that is being scanned.

The last category (L0H3, L1H1, L2H0, L1H5, L2H1, L2H5, L4H5) is also located near the main diagonal but displays broader diagonals, reflecting attention distributed over a wider local window. These heads indicate that the model integrates information from a broader local sequence context around each position, suggesting that codon choice depends on short-range contextual information spanning several neighboring codons.

Attention peaks are observed either symmetrically around the main diagonal or predominantly in one direction within a given head (upstream or downstream). In some cases, unidirectional heads have a complementary head showing an approximately symmetric pattern (e.g., L2H2 and L2H3, or L2H6 and L1H4). This suggests that unidirectional heads focus on specific motifs or codons, consistent with the first category, where attention is concentrated on very precise positions. In the second category, heads tend to capture information either upstream or downstream. By contrast, the broader and more symmetric diagonals in the last category likely reflect the overall local frequency of codons around a position, which naturally leads to symmetric attention. Across categories, attention is often slightly biased toward downstream positions relative to the predicted codon (Fig.5 C-D).

### 2.5 Prediction of codon-level functional constraints in experimental datasets

Although synonymous mutations are not completely neutral as initially assumed, they do not all have the same impact, nor do they all significantly affect protein function or fitness^39^. Some positions are extremely constrained and tolerate only the wild-type codon, others show relaxed constraints and tolerate only a subset of synonymous codons, and finally some positions accept all synonymous codons as interchangeable. Positions under strong codon-level selective pressure are therefore particularly important for studying the mechanisms underlying codon bias.

Sequence databases contain only wild-type codon sequences, and codon prediction models are trained and evaluated on recovery of the natural codon, without considering whether individual positions are under strong or weak synonymous constraint. Because such functional information is rarely available, it cannot currently be incorporated at scale.

To evaluate the true ability of CaNAT to account for codon-bias constraints at individual positions, we applied the model to three experimental datasets measuring the effects of systematic synonymous mutations on protein fitness. All datasets consist of *E. coli* proteins expressed in *E. coli*, ensuring that native codon-usage constraints are preserved.

D-alanine—D-alanine ligase A (ddlA) is a native protein of E.coli, catalyzing the formation of D-alanine dipeptides. ddlA is a non-essential protein with low capability to refold in the absence of chaperones, making it a interesting system for investigating how co-translational folding through the ribosome is affected by codon usage. Folding efficiency of ddlA was investigated with deep synonymous scanning in^40^. In this study, 22 positions with complete synonymous coverage were analyzed using protein folding efficiency assay, which we use here as a proxy for fitness. *RNase III* is a native, non-essential E. coli protein that cleaves double-stranded RNA and plays a key role in RNA processing and regulation. *RNase III* activity was investigate with DSS^41^. A functional score was used to quantify how well each mutation preserves RNase III activity, based on its ability to repress GFP in a reporter system. In our study the 222 positions were analyzed using the functional score to decide of mutation consequences. *TEM1 β-lactamase* is a hydrolase responsible for antibiotic resistance and a commonly found as an plasmid-encoded protein in E. coli, present since the 1960s^42^, and has since been exposed to selection pressures associated with E. coli codon usage. 272 positions with full coverage were included in our analysis. Methionine and tryptophan were excluded from all analyses.

Each position was classified into one of three categories based on experimental fitness measurements across all synonymous codons (see Methods). Positions were labeled WO (wild-type only tolerated) when only the native codon maintained fitness, PS (partially synonymous tolerated) when only a subset of synonymous codons was compatible with normal fitness, and FS (fully synonymous tolerated) when all synonymous codons were functionally equivalent. This classification allows direct assessment of model performance as a function of the strength of codon-level selective constraints at each position.

For each category (WO, PS, FS), two performance metrics were evaluated: the accuracy in recovering the wild-type codon and the accuracy in predicting any non-deleterious codon (Figure 5 A,C,E). In addition, the proportion of wild-type rare codons in each category was analyzed (Fig. 2.5 B,D,F). Prediction accuracy for the wild-type codon was highest in the WO category compared to PS and FS positions. Notably, at PS positions, where several but not all synonymous codons were tolerated, CaNAT often failed to recover the exact wild-type codon but frequently predicted another tolerated variant. This pattern suggests that the model does not simply mimics wild-type codon usage, but may also reflect the partial relaxation of synonymous constraints at these sites.

Importantly, the WO category contained a higher proportion of rare codons across all datasets, which are generally more difficult to predict. Despite this, CaNAT achieved its best performance in this most constrained category, indicating that the model is capable of detecting strong codon-level selective signals, particularly at positions where only a single, often rare, codon is functionally tolerated.

## 3 Discussion

In this study, we developed a Transformer-based model, CaNAT, designed to predict synonymous codon usage along coding sequences. Unlike previous models, CaNAT accurately captures rare codons and provides a reliable confidence score that quantifies prediction uncertainty. Although the model does not outperform existing methods for all codons, its interpretability and capacity to recover biologically relevant biases represent a major step toward context-aware codon prediction.

Our analysis shows that CANAT implicitly learns species-specific codon usage biases, even in the absence of explicit annotation during training. The patterns captured by the model are consistent with known biological preferences, indicating that amino-acid sequences alone provide sufficient information to recover both species identity and characteristic codon usage, while preserving context-dependent nuances in codon selection.

Importantly, the task of synonymous codon prediction appears to constrain the model to attend to subtle, context-dependent features of protein sequences that are informative of the host organism. This constraint likely promotes the emergence of species-discriminative representations, enabling accurate classification of individual positions according to their corresponding species (Tab. 2). Together, these results suggest that codon prediction constitutes a biologically meaningful task that naturally encourages a model to encode organism-level information beyond primary sequence identity.

**Table 2:** Confusion matrix of LDA classifier between E coli, H sapiens ans S thermophilus based on embeddings.

| True / Predicted | Homo sapiens | Escherichia coli | Streptococcus thermophilus |
| --- | --- | --- | --- |
| **Homo sapiens** | 30104 | 826 | 153 |
| **Escherichia coli** | 904 | 20439 | 3066 |
| **Streptococcus thermophilus** | 433 | 1582 | 21444 |

Independently, CaNAT also appears to capture features related to RNA secondary structure and stability, suggesting that it encodes functionally meaningful dependencies between sequence context and codon choice. The structured attention patterns uncovered here suggest that the model captures biologically meaningful dependencies related to the translation process. Short-range diagonal motifs likely reflect local constraints on codon choice, potentially associated with interactions between neighboring codons, such as effects on tRNA availability, ribosomal decoding kinetics, or nascent peptide folding at the ribosomal exit tunnel. These windowed patterns are consistent with the concept of dicodons. The strongest signal comes from directly adjacent codon pairs, but non-adjacent codon pairs also carry informative signals, albeit with smaller magnitude. This observation extends the dicodon notion toward broader, short-range contextual influences, in line with the work of Gutman et al. ^33^, who reported that in Escherichia coli, codon pairs separated by two or three intervening codons still exhibit non-random usage, even if substantially weaker than adjacent biases. By leveraging a large sequence corpus and the ability of transformer models to detect subtle but consistent patterns, our results provide further support for this extension of dicodon bias to a slightly broader window. The extension of these motifs beyond immediately adjacent codons is also consistent with experimental evidence that translational dynamics can be influenced by short stretches of consecutive codons rather than isolated positions.

Broader local attention windows may correspond to regulatory features acting over extended local contexts, such as modulation of elongation rates, avoidance of translational pausing, or maintenance of reading frame stability. These patterns suggest that synonymous codon selection is not determined solely by individual codon optimality but also by the surrounding sequence context.

Long-range attention patterns are particularly intriguing, as they imply that distant regions of the coding sequence contribute to local codon decisions. Such dependencies may reflect higher-order translational constraints, including coordination between early and late elongation phases, co-translational folding of protein domains, or global regulation of translation efficiency across the coding sequence. The observed asymmetry favoring downstream positions is consistent with a causal interpretation in which future sequence context informs current codon choice, potentially reflecting anticipation of upcoming translational constraints.

Together, these observations suggest that the model does not rely solely on global gene-level features, such as overall expression level or codon usage bias, but instead integrates multi-scale contextual information that is compatible with known principles of translational regulation and link with its ability to perceived functional protein pressure. An additional consequence of being sensitive to multiscale constraints is that CaNAT effectively identifies positions under strong codon-level selection. Across the sequence, it tends to predict codons that preserve fitness: the wild-type codon at positions under strict selective constraint, and one of several possible codons at more tolerant positions. This suggests that the model indirectly captures the influence of selection pressure, partially reflecting fitness effects. The patterns of codon choice inferred by the model likely emerge from the large corpus of amino acid sequences used for training, highlighting underlying functional or structural constraints on proteins and their relationship with codon usage bias. These results suggest a novel approach for investigating how codon usage may influence protein structure and function. While correlations with secondary structure have been reported previously, the predicted codon patterns here hint that it is possible to train models to capture aspects of sequence-dependent fitness, opening new avenues for exploring genotype-to-phenotype relationships.

Taken together, these results illustrate that a Transformer-based model such as CaNAT is particularly well-suited to integrate multiple regulatory layers, from RNA structure to translational dynamics, within a unified representation, offering a powerful framework to study the complexity of codon usage. Overall, CaNAT provides a robust framework for detecting and characterizing rare codons in coding sequences, offering new insights into how synonymous choice shapes translational dynamics. By capturing context-dependent patterns of codon usage, it highlights regions where codon composition may influence expression levels, folding kinetics, or protein stability. Beyond variant interpretation, these findings open perspectives for rational gene design, for instance, optimizing heterologous expression, fine-tuning translation rates, or correcting deleterious synonymous patterns in therapeutic constructs. In the longer term, integrating CaNAT with models of mRNA dynamics or co-translational folding could contribute to predictive, sequence-level control of protein expression across organisms.

## 4 Methods

### Datasets

Coding DNA sequences (CDS) were downloaded on September 8, 2023 from the European Nucleotide Archive (ENA) coding sequence repository (ftp://ftp.ebi.ac.uk/pub/databases/ena/coding/). CDS were retained only if they started with the canonical start codon (ATG) and contained exclusively valid nucleotides (A, T, G, and C). Sequences were translated into amino acid sequences and used for all subsequent clustering steps.

The protein dataset was clustered at 30% identity and 90% coverage using MMseqs2 (version 14-7e284) with the command `linclust –min-seq-id 0.3 -c 0.9`. To prevent homology leakage between datasets, entire clusters were assigned exclusively to either the training, validation, or test set, ensuring that no pair of sequences across different sets exceeded 30% sequence identity.

Each dataset was constrained to contain sequences from at least three model species: *Escherichia coli, Homo sapiens*, and *Saccharomyces cerevisiae*. The validation and test sets were required to include sequences from at least 100 distinct species and to contain no more than 200,000 sequences in order to ensure sufficient diversity for evaluation while preserving an adequate number of sequences for training. To further reduce redundancy within species, a second clustering step was performed independently for each species using MMseqs2 `linclust` with a minimum sequence identity threshold of 0.90 and a minimum alignment coverage of 0.50. Only the representative sequence of each cluster was retained. After this second clustering step, the final datasets used for model training, test and evaluation comprised approximately 3 million sequences in the training set, 500 sequences in the validation set, and 200 sequences in the test set. Redundancy across species was intentionally preserved to allow the model to capture species-specific codon usage patterns.

### Architecture and Training process

Protein sequences were tokenized using the amino acid to token dictionary of ProteinMPNN^43^. Tokens were embedded into a 512-dimensional latent space and combined with positional encodings.

The model was implemented in PyTorch using a Transformer encoder/decoder architecture. The encoder consisted of 6 layers with 8 attention heads per layer and a model dimension of 512, and produced contextual representations of the amino acid sequence. The decoder also consisted of 6 layers with identical dimensionality and integrated both the encoder outputs through cross-attention and the embedded amino acid sequence as input. No causal masking was applied in the decoder to enable fully parallel prediction across positions and improve computational efficiency. Preliminary experiments did not indicate a consistent performance benefit from masking for this task.

The decoder outputs were projected to codon logits using a fully connected linear layer, and the model was trained to directly predict the corresponding codon at each amino acid position.

Sequences were padded or truncated to a common length of 512 codons and trained with a batch size of 64. Optimization was performed using the Adam optimizer with a learning rate of 1 *×* 10^−4^. The training objective was a weighted cross-entropy loss, where codon frequencies were computed within each batch and used to compensate for class imbalance.

Training was initialized with 100 optimization steps using single synthetic random sequences, allowing the model to first learn the basic genetic code mapping.

### Model confidence evaluation

CaNAT outputs a score for each codon at every sequence position. Confidence scores were computed by applying a softmax over all codons at each position, yielding values between 0 and 1. Because predicted probabilities are mostly distributed among synonymous codons, absolute confidence varies with the number of synonyms (1–6), complicating direct comparisons. To account for this, we computed a normalized confidence measure described (Fig. 1 F).

### Rare codon assignment

For each amino acid, codon frequencies were normalized across the full set of synonymous codons using reference frequencies from the Kazuka tables (^31^ downloaded 08/2025) to calculate the RSCU for each codon. Codons with RSCU < 0.7 were considered rare.

### Statistical analysis

#### Spearman correlation

For each species (*Streptococcus thermophilus, Homo sapiens*, and *Escherichia coli* the frequencies of every codon by amino acid and the predicted one have been recorded over the training set (for positions 0 to 512 in sequences), and Spearman correlation have been calculated by `scikit-learn` by AA and distribution of AA spearman serve to construct general boxplot.

#### LDA

For each species (*Streptococcus thermophilus, Homo sapiens*, and *Escherichia coli*), codon embeddings extracted of the decoder module just before the final projection layer of 500 random sequences (for positions 0 to 512 in sequences) were recorded. A Linear Discriminant Analysis (`n_components=2`, `scikit-learn`) was trained to separate species on embeddings of 500 random sequences, and embeddings of 100 distinct sequences per species from a distinct test set were projected onto the LDA space. Two-dimensional kernel density estimates of the test embeddings were visualized with five contour levels using `seaborn.kdeplot`, colored by species.

#### Stability

RNA secondary structure stability was calculated using the `ViennaRNA` package (version 2.7.0) ^44^

#### Regression Stability-Accuracy accounting for GC-content

Two linear models were then constructed to explain accuracy: one with GC-content alone, and another with GC-content and predicted stability as explanatory variables. Both models were fitted using the Ordinary Least Squares (OLS) implementation from the Python `statsmodels` module, which was also used to extract corresponding *R*^2^ values and *p*-values. For relevant statistical analyses, a random subset of 10,000 sequences from the training set was used.

Partial regression plots were used to visualize the effect of stability on accuracy after controlling for GC-content. Specifically, accuracy and stability were each regressed on GC-content using ordinary least squares, and the residuals from these regressions were plotted against each other.

#### Attention map

Attention maps were extracted during inference for all sequences in the test set, across all attention layers and heads. A subset of layers and heads exhibiting a clear diagonal pattern was subsequently selected based on visual inspection. Sequences longer than 512 codons were truncated to a maximum length of 512, while shorter sequences were retained at their original length without padding. every attention head have been normalized by `minMax` operation. To compute average attention maps, attention weights were aggregated independently for each position pair (i,j) by averaging only over sequences in which both positions were defined. For visualization (Fig. 5 A–B), attention maps were independently min–max normalized for each layer–head pair and overlaid as heatmaps. Superimposed attention patterns were further highlighted using contour representations. Attention scores were normalized within each head and pooled across heads. An attention-weighted histogram of positional offsets was computed within a restricted central range to characterize the distribution of relative attention across offsets.

### Mutational scanning datasets

The codon-level mutational dataset of ddlA was obtained from^40^. The reported normalized fluorescence ratio (norm RFP/GFP) was used as a proxy for codon tolerance. Only codon positions for which all synonymous codons were experimentally covered were retained for analysis. A threshold of 1 was used to consider a codon as tolerated (>1) or not (<1), with all wild-type (WT) codons having an exact value of 1.

Codon mutation data for RNase III were extracted from^41^, Supplementary Data S8. The reported functional score was used as a proxy for codon mutation tolerance. Only synonymous substitutions were retained. functional score and ‘weighted mean’ between replicas. A threshold of 0 was used to consider a codon as tolerated (>0) or not (<0), with all wild-type (WT) codons having an exact value of 0.

Codon mutational scanning data for TEM-1 *β*-lactamase were obtained from^45^, Supplementary file `supp_msu081_Data_S1–S4.xlsx`. The reported fitness values were used as measures of codon tolerance. Only synonymous mutations were retained. A threshold of 1 was used to consider a codon as tolerated (>1) or not (<1), with all wild-type (WT) codons having an exact value of 1.

For all datasets, the corresponding wild-type protein sequences were used as input to CaNAT to generate codon-level predictions, which were then evaluated against experimentally measured effects of synonymous codon substitutions.

## Code availability

Model architecture, parameters, and additional codes are available at https://github.com/Andre-lab/CaNAT/.

## References

[1] Plotkin, J. B. & Kudla, G. Synonymous but not the same: the causes and consequences of codon bias. Nature Reviews Genetics 12, 32–42 (2011).

[2] Chamary, J.-V., Parmley, J. L. & Hurst, L. D. Hearing silence: non-neutral evolution at synonymous sites in mammals. Nature Reviews Genetics 7, 98–108 (2006).

[3] Shabalina, S. A., Spiridonov, N. A. & Kashina, A. Sounds of silence: synonymous nucleotides as a key to biological regulation and complexity. Nucleic acids research 41, 2073–2094 (2013).

[4] Tchebotarev, L. & Herzel, L. Secret code: Encoding promoters by synonymous codons. Proceedings of the National Academy of Sciences 121, e2416360121 (2024).

[5] Radrizzani, S., Kudla, G., Izsvák, Z. & Hurst, L. D. Selection on synonymous sites: the unwanted transcript hypothesis. Nature Reviews Genetics 25, 431–448 (2024).

[6] Sarkar, A., Panati, K. & Narala, V. R. Code inside the codon: The role of synonymous mutations in regulating splicing machinery and its impact on disease. Mutation Research/Reviews in Mutation Research 790, 108444 (2022).

[7] Parmley, J. L. & Hurst, L. D. Exonic splicing regulatory elements skew synonymous codon usage near intron-exon boundaries in mammals. Molecular biology and evolution 24, 1600–1603 (2007).

[8] Wang, Y., Qiu, C. & Cui, Q. A large-scale analysis of the relationship of synonymous snps changing microrna regulation with functionality and disease. International journal of molecular sciences 16, 23545–23555 (2015).

[9] Sauna, Z. E. & Kimchi-Sarfaty, C. Understanding the contribution of synonymous mutations to human disease. Nature Reviews Genetics 12, 683–691 (2011).

[10] Cartegni, L., Chew, S. L. & Krainer, A. R. Listening to silence and understanding nonsense: exonic mutations that affect splicing. Nature reviews genetics 3, 285–298 (2002).

[11] Liu, Y., Yang, Q. & Zhao, F. Synonymous but not silent: the codon usage code for gene expression and protein folding. Annual review of biochemistry 90, 375–401 (2021).

[12] Komar, A. Synonymous codon usage—a guide for co-translational protein folding in the cell. Molecular Biology 53, 777–790 (2019).

[13] Parvathy, S. T., Udayasuriyan, V. & Bhadana, V. Codon usage bias. Molecular biology reports 49, 539–565 (2022).

[14] Kames, J. et al. Tissuecocoputs: novel human tissue-specific codon and codon-pair usage tables based on differential tissue gene expression. Journal of molecular biology 432, 3369–3378 (2020).

[15] Liu, Y. A code within the genetic code: codon usage regulates co-translational protein folding. Cell Communication and Signaling 18, 145 (2020).

[16] Zhao, F., Yu, C.-h. & Liu, Y. Codon usage regulates protein structure and function by affecting translation elongation speed in drosophila cells. Nucleic acids research 45, 8484–8492 (2017).

[17] Dilucca, M., Cimini, G., Forcelloni, S. & Giansanti, A. Co-evolution between codon usage and protein-protein interaction in bacteria. Gene 778, 145475 (2021).

[18] Moss, M. J., Chamness, L. M. & Clark, P. L. The effects of codon usage on protein structure and folding. Annual Review of Biophysics 53 (2024).

[19] Chaney, J. L. & Clark, P. L. Roles for synonymous codon usage in protein biogenesis. Annual review of biophysics 44, 143–166 (2015).

[20] Sharp, P. M. & Li, W.-H. The codon adaptation index-a measure of directional synonymous codon usage bias, and its potential applications. Nucleic acids research 15, 1281–1295 (1987).

[21] Sabi, R., Volvovitch Daniel, R. & Tuller, T. staicalc: trna adaptation index calculator based on species-specific weights. Bioinformatics 33, 589–591 (2017).

[22] Rodriguez, A., Wright, G., Emrich, S. & Clark, P. L. % minmax: a versatile tool for calculating and comparing synonymous codon usage and its impact on protein folding. Protein Science 27, 356–362 (2018).

[23] Orešič, M. & Shalloway, D. Specific correlations between relative synonymous codon usage and protein secondary structure. Journal of molecular biology 281, 31–48 (1998).

[24] Sidi, T., Bahiri-Elitzur, S., Tuller, T. & Kolodny, R. Predicting gene sequences with ai to study codon usage patterns. Proceedings of the National Academy of Sciences 122, e2410003121 (2025).

[25] Jain, R., Jain, A., Mauro, E., LeShane, K. & Densmore, D. Icor: improving codon optimization with recurrent neural networks. BMC bioinformatics 24, 132 (2023).

[26] Outeiral, C. & Deane, C. M. Codon language embeddings provide strong signals for use in protein engineering. Nature Machine Intelligence 6, 170–179 (2024).

[27] Fallahpour, A., Gureghian, V., Filion, G. J., Lindner, A. B. & Pandi, A. Codontransformer: a multispecies codon optimizer using context-aware neural networks. Nature Communications 16, 3205 (2025).

[28] Han, X. et al. Deepcodon: A deep learning codon-optimization model to enhance protein expression. BioDesign Research 100042 (2025).

[29] Fu, H. et al. Codon optimization with deep learning to enhance protein expression. Scientific reports 10, 17617 (2020).

[30] Chowdhury, T., Saha, A., Saha, A., Chakraborty, A. & Das, N. Neuralcodopt: Codon optimization for the development of dna vaccines. Computational Biology and Chemistry 116, 108377 (2025).

[31] Nakamura, Y., Gojobori, T. & Ikemura, T. Codon usage tabulated from international dna sequence databases: status for the year 2000. Nucleic acids research 28, 292–292 (2000).

[32] Frumkin, I. et al. Codon usage of highly expressed genes affects proteome-wide translation efficiency. Proceedings of the National Academy of Sciences 115, E4940–E4949 (2018).

[33] Gutman, G. A. & Hatfield, G. W. Nonrandom utilization of codon pairs in escherichia coli. Proceedings of the National Academy of Sciences 86, 3699–3703 (1989).

[34] Tats, A., Tenson, T. & Remm, M. Preferred and avoided codon pairs in three domains of life. BMC genomics 9, 463 (2008).

[35] Kunec, D. & Osterrieder, N. Codon pair bias is a direct consequence of dinucleotide bias. Cell reports 14, 55–67 (2016).

[36] Irwin, B., Heck, J. D. & Hatfield, G. W. Codon pair utilization biases influence translational elongation step times (). Journal of Biological Chemistry 270, 22801–22806 (1995).

[37] Gamble, C. E., Brule, C. E., Dean, K. M., Fields, S. & Grayhack, E. J. Adjacent codons act in concert to modulate translation efficiency in yeast. Cell 166, 679–690 (2016).

[38] Alonso, A. M. & Diambra, L. Dicodon-based measures for modeling gene expression. Bioinformatics 39, btad380 (2023).

[39] Love, A. M. & Nair, N. U. Specific codons control cellular resources and fitness. Science Advances 10, eadk3485 (2024).

[40] Otsuka, F. A. & André, I. Effects of single synonymous substitutions on folding efficiency demonstrate the influence of rare codons and protein structure. bioRxiv 2025–03 (2025).

[41] Weeks, R. & Ostermeier, M. Fitness and functional landscapes of the e. coli rnase iii gene rnc. Molecular Biology and Evolution 40, msad047 (2023).

[42] Bush, K. Past and present perspectives on β-lactamases. Antimicrobial agents and chemotherapy 62, 10–1128 (2018).

[43] Dauparas, J. et al. Robust deep learning–based protein sequence design using proteinmpnn. Science 378, 49–56 (2022).

[44] Lorenz, R. et al. Viennarna package 2.0. Algorithms for molecular biology 6, 26 (2011).

[45] Firnberg, E., Labonte, J. W., Gray, J. J. & Ostermeier, M. A comprehensive, high-resolution map of a gene’s fitness landscape. Molecular biology and evolution 31, 1581–1592 (2014).

