## Supplementary material for "Decoding synonymous codon selection with a Transformer model": Supp figures

### Supplementary Figures

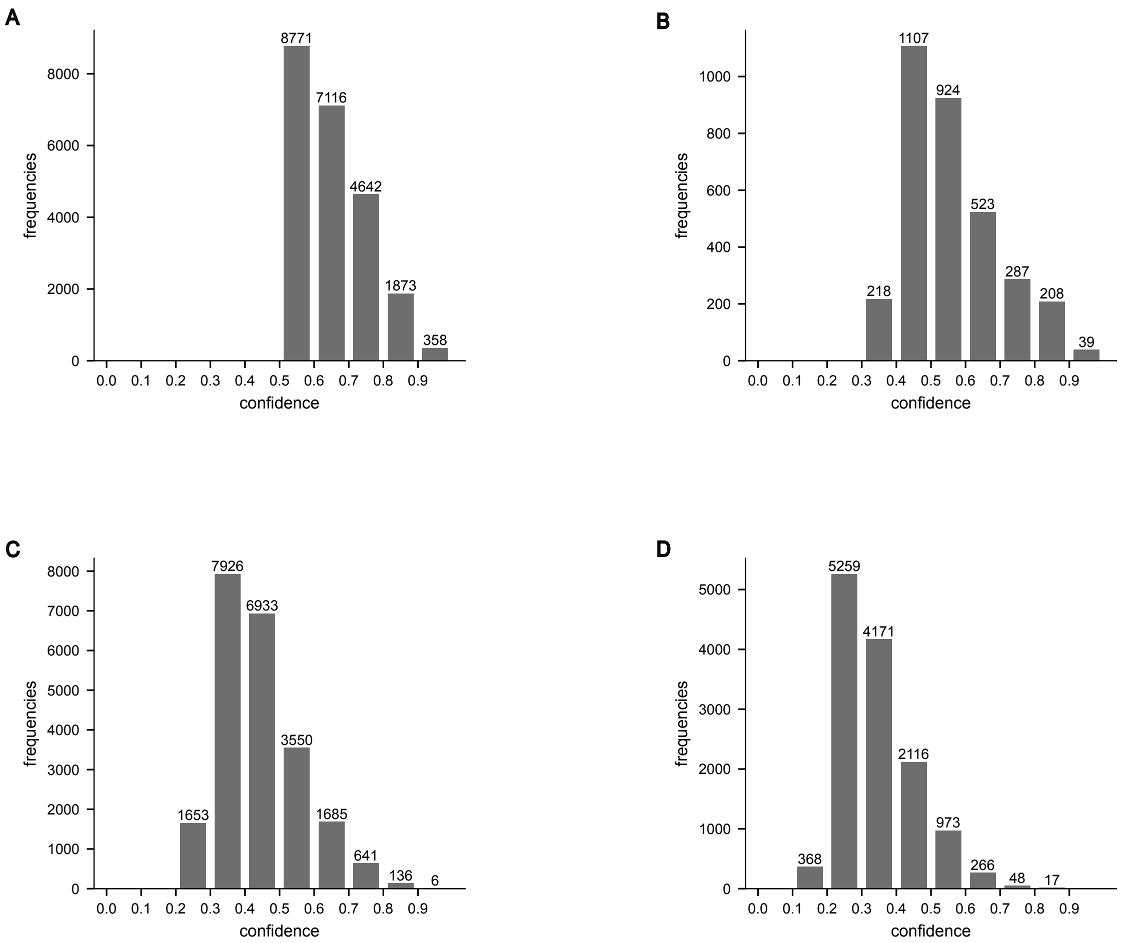


Figure S1: Distribution of model confidence scores for codon predictions in the test set. (A) Amino acids encoded by 2 synonymous codons. (B) Amino acids encoded by 3 synonymous codons. (C) Amino acids encoded by 4 synonymous codons. (D) Amino acids encoded by 6 synonymous codons.

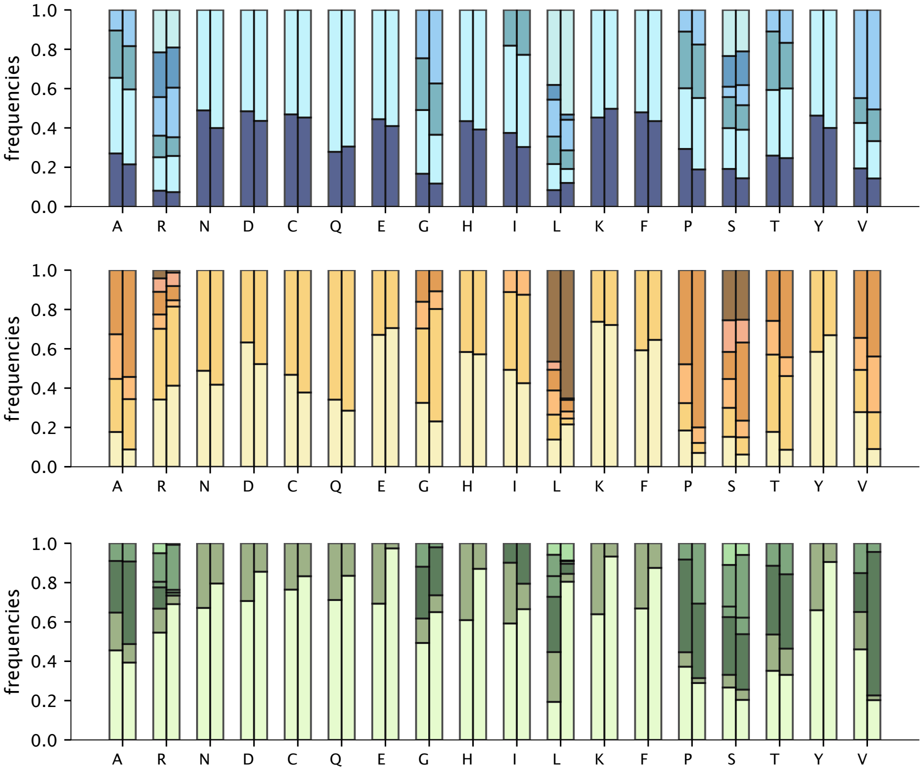


Figure S2: Stacked bar plots of normalized synonymous codon frequencies by amino acid, comparing natural (left) and predicted (right) distributions. Upper plot: *Homo sapiens* (blue); middle plot: *Escherichia coli* (orange); lower plot: *Streptococcus thermophilus* (green).

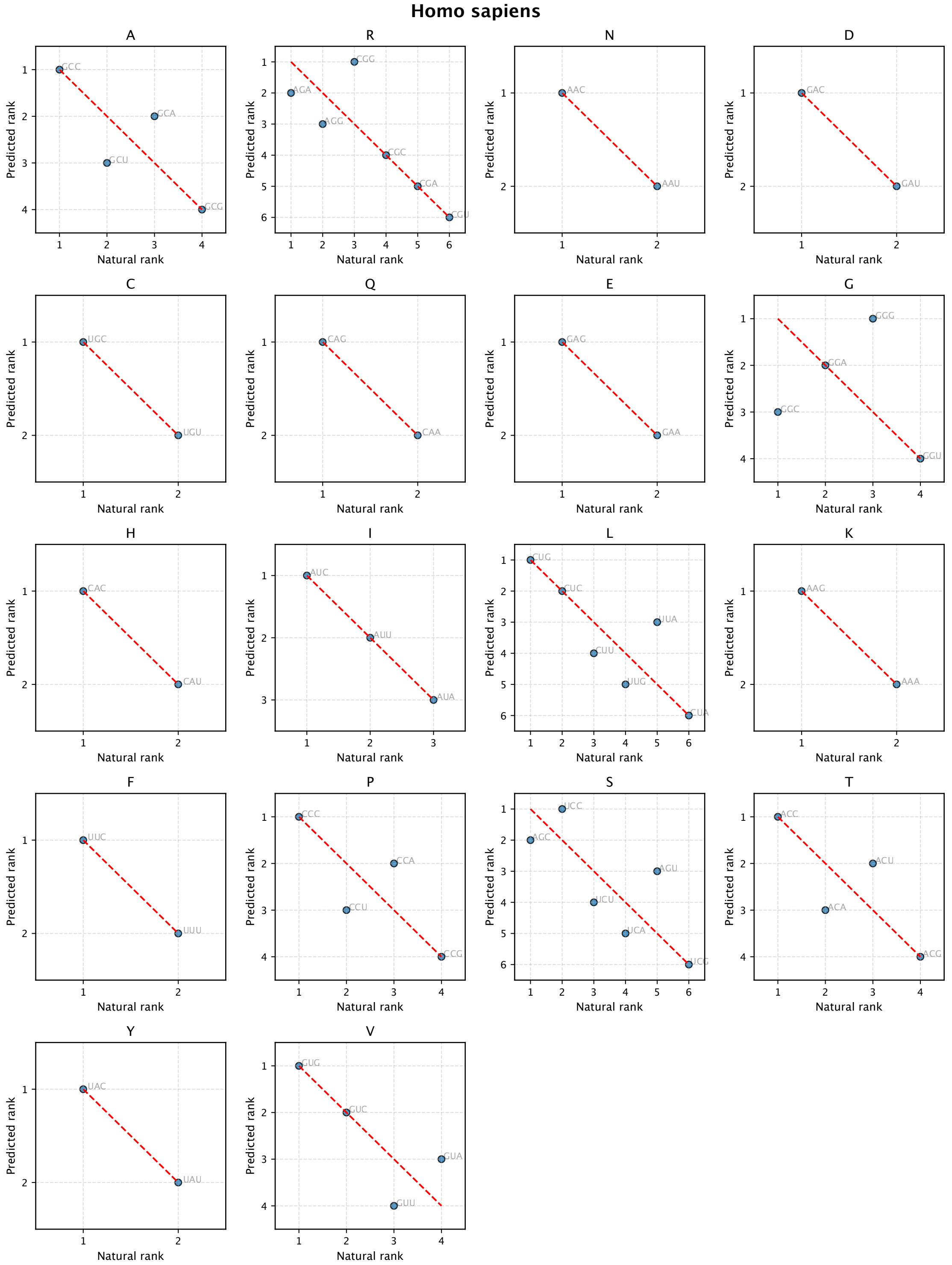


Figure S3: Rank–rank plots showing the correspondence between natural and predicted synonymous codon ranks for all 20 amino acids, one panel per amino acid, for *Homo sapien*.

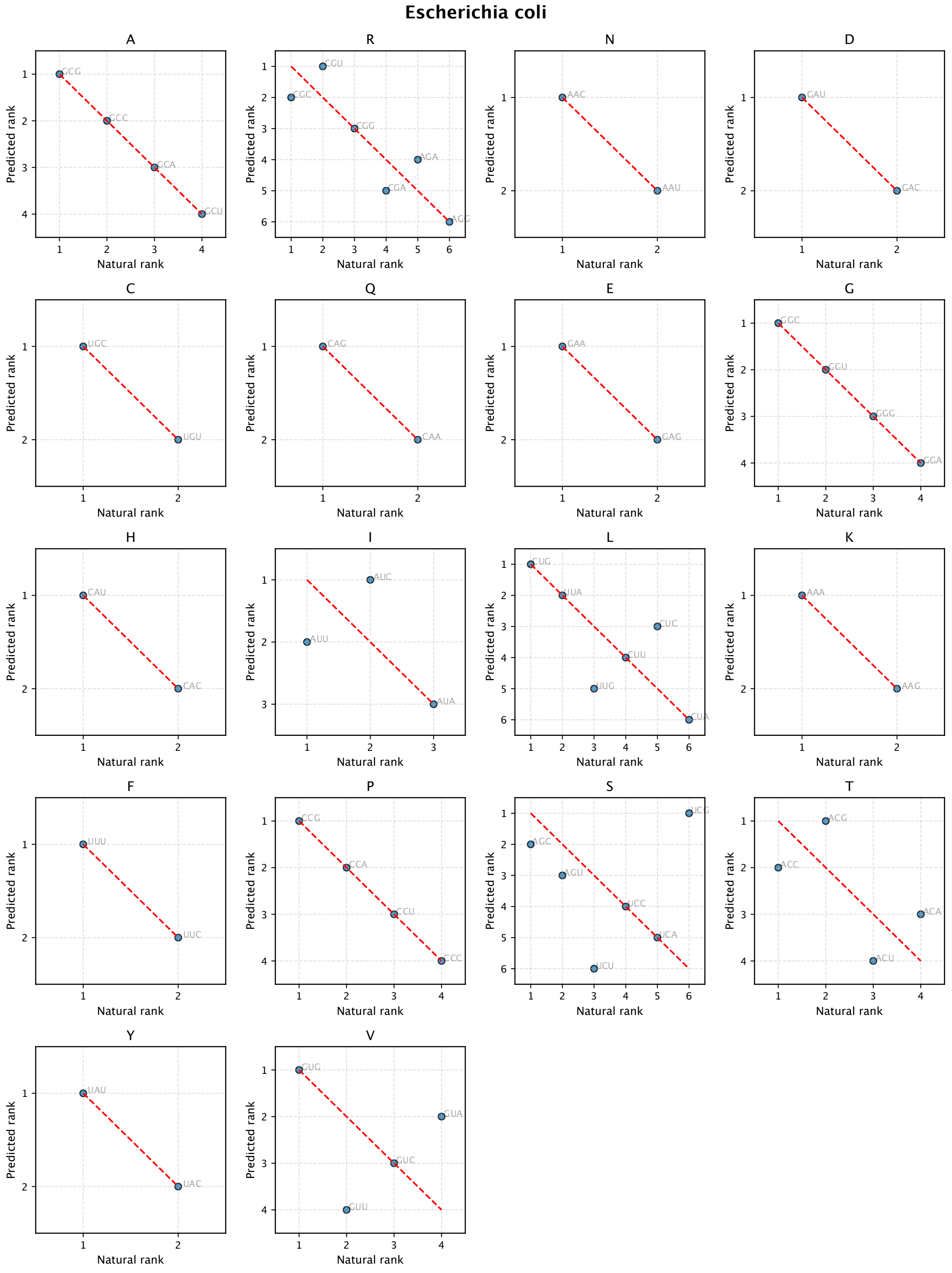


Figure S4: Rank–rank plots showing the correspondence between natural and predicted synonymous codon ranks for all 20 amino acids, one panel per amino acid, for *Escherichia coli*.

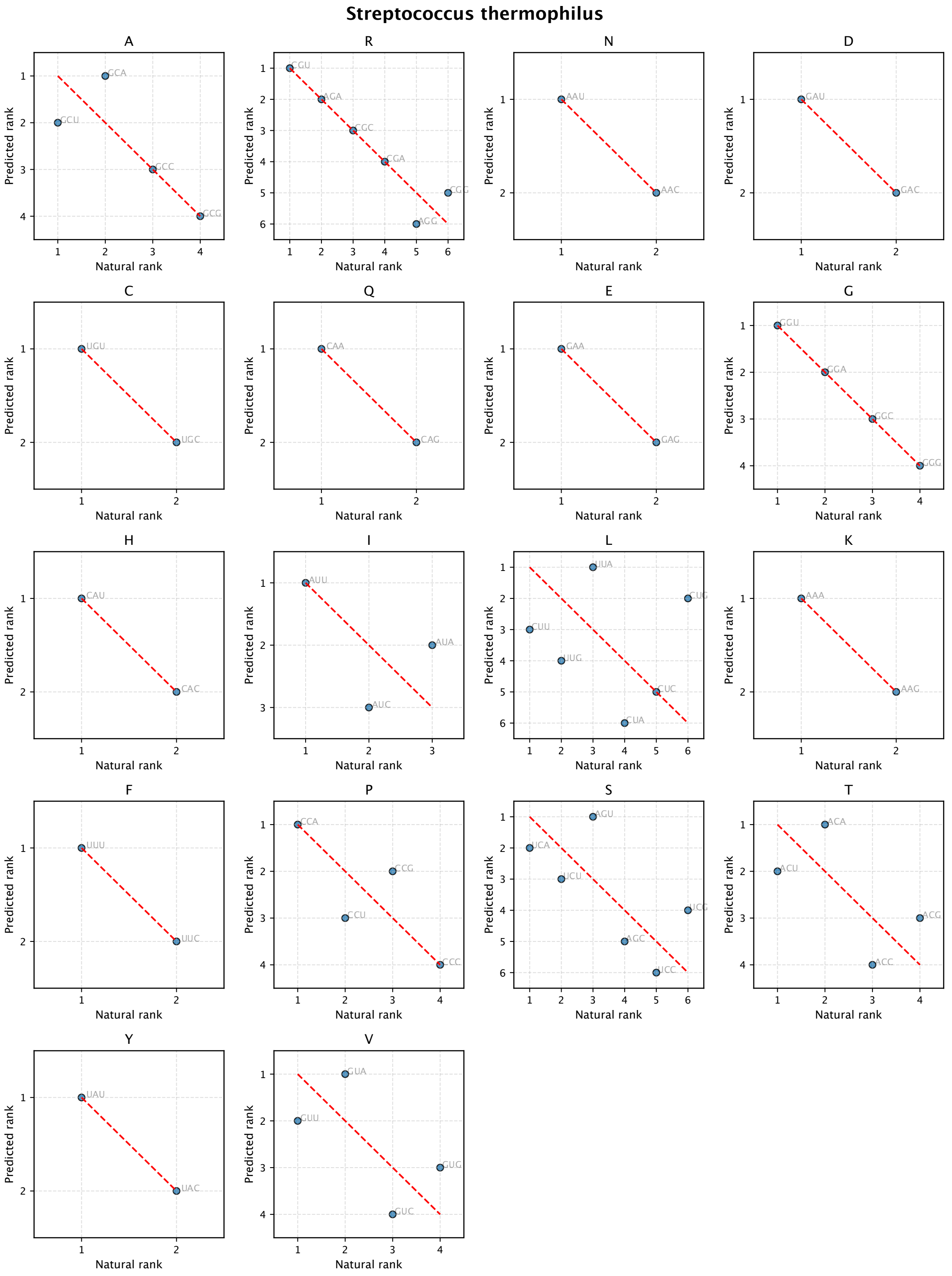


Figure S5: Rank–rank plots showing the correspondence between natural and predicted synonymous codon ranks for all 20 amino acids, one panel per amino acid, for *Streptococcus thermophilus*.

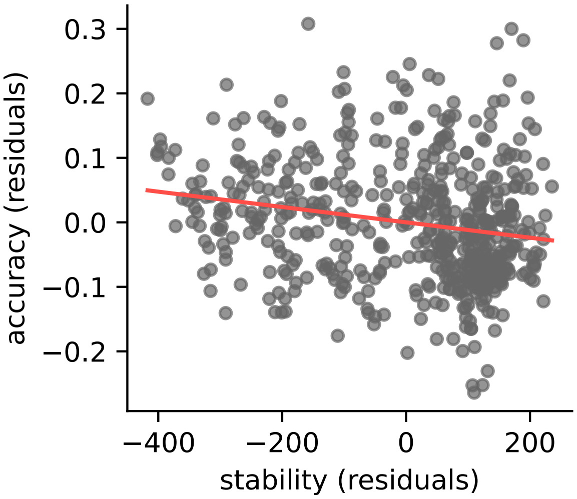


Figure S6: **Partial regression plot** of residual accuracy versus residual stability after controlling for GC-content. The line indicates the regression slope ($-1.19 \times 10^{-4}$). The partial correlation, quantifying the strength of this association conditional on GC-content, is $-0.20$

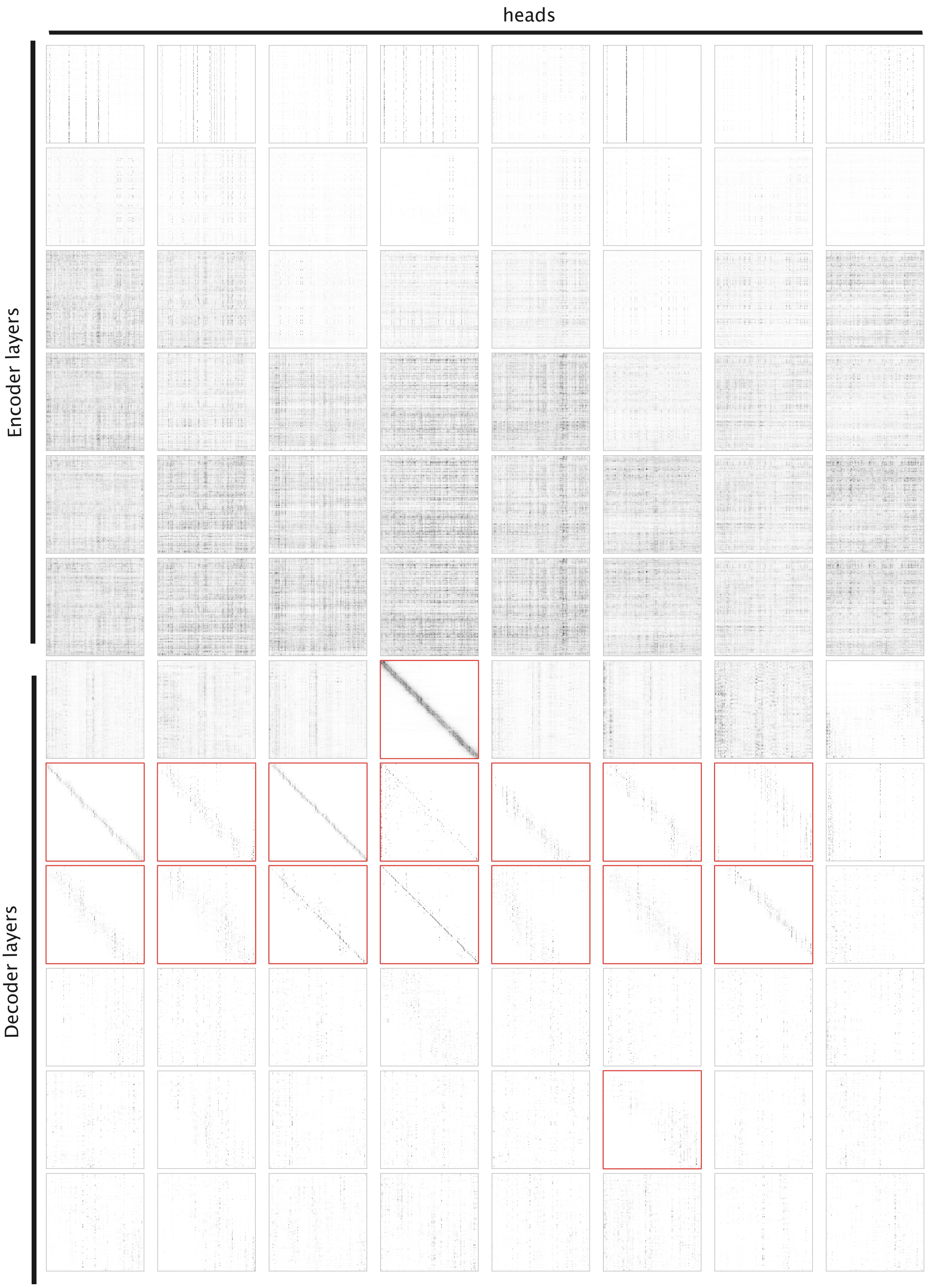


Figure S7: All 8 attention heads from each of the 6 encoder and 6 decoder layers are shown for the example protein ENA—KWV17632.1 (*Escherichia coli* oxidative-stress-resistance chaperone). Heads retained for diagonal analysis are highlighted with a red border.
